# Uncovering Medical Insights from Vast Amounts of Biomedical Data in Clinical Case Reports

**DOI:** 10.1101/172460

**Authors:** Yijiang Zhou, David A. Liem, Jessica M. Lee, Quan Cao, Brian Bleakley, J. Harry Caufield, Sanjana Murali, Wei Wang, Li Zhang, Alex Bui, Yizhou Sun, Karol E. Watson, Jiawei Han, Peipei Ping

## Abstract

Clinical case reports (CCRs) have a time-honored tradition in serving as an important means of sharing clinical experiences on patients presenting with atypical disease phenotypes or receiving new therapies. However, the huge amount of accumulated case reports are isolated, unstructured, and heterogeneous clinical data, posing a great challenge to clinicians and researchers in mining relevant information through existing indexing tools. In this investigation, in order to render CCRs more findable, accessible, interoperable, and reusable (FAIR) by the biomedical community, we created a resource platform, including the construction of a test dataset consisting of 1000 CCRs spanning 14 disease phenotypes, a standardized metadata template and metrics, and a set of computational tools to automatically retrieve relevant medical information and to analyze all published PubMed clinical case reports with respect to trends in publication journals, citations impact, MeSH Terms, drug use, distributions of patient demographics, and relationships with other case reports and databases. Our standardized metadata template and CCR test dataset may be valuable resources to advance medical science and improve patient care for researchers who are using machine learning approaches with a high-quality dataset to train and validate their algorithms. In the future, our analytical tools may be applied towards other large clinical data sources as well.

## Introduction

Clinical case reports serve as an important means of sharing observations, discussions, and novel insights about patient presentations with unique disease phenotypes or new therapies, making them an invaluable source for patient care, scientific research, and epidemiologic investigations.[1] Throughout history, case reports have traditionally represented the initial account of a rare disease, often with its genetic background.[2] Two important examples that highlight the pivotal impact of reporting new clinical observations are the first treatment of human rabies by Louis Pasteur in 1885 and the first application of penicillin in patients in 1943.[3, 4] Moreover, presenting the case report as a coherent narrative that evolves over time allows the reader to contemplate the disease course as well as the diagnostic and therapeutic reasoning.

Over the past decades, medical data and clinical information have continued to accumulate through clinical case reports, most of which can be accessed through the PubMed Database.[5] As of July 2017, a total of ∼1.8 million case reports can be retrieved using the PubMed search engine, with half a million published in the past decade alone. The data from available case reports, as well as the data describing case report data (metadata), may contain unexplored knowledge that could prompt new investigations and facilitate the characterization of individual patients in precision medicine.[6-9] Unfortunately, due to the staggering size of the case report corpus, the complexity and unstructured state of case report data, and heterogeneity in case report content, efforts to thoroughly process case report data remains challenging. Nonetheless, the development of new methods to better index, annotate, and curate case reports will become essential to extracting relevant medical information. In this manuscript, we devised a resource platform, including the construction of a test data set consisting of 1000 CCRs spanning 14 disease phenotypes and standardization of a metadata template and metrics. We further established a set of computational tools to automatically retrieve relevant medical information and to analyze the available online clinical case reports with respect to trends in publication journals, citations impact, MeSH Terms, drug use, distributions of patient demographics, and relationships with other CCRs and databases.

### Merits and current challenges of clinical case report publications

By December 2016, the number of case report publications on PubMed under the MeSH term of “Diseases Category” totaled to 1,724,755, with cancer, neurological disease, and cardiovascular disease as the three most studied and published disease areas reported (Figure 1). The total number of published case reports has been increasing over the past 40 years (1975-2015), though their growth rate has been slower than that of the total accumulated publications (Figure 2A). A similar trend was also observed in three prominent journals: New England Journal of Medicine, Lancet, and Journal of American Medical Association (Figure 2A). This may be explained by the scarcity of rare disease cases; alternatively, the relatively low citation rate of case reports may make case reports less appealing to journals.[10]

**Figure 1:**
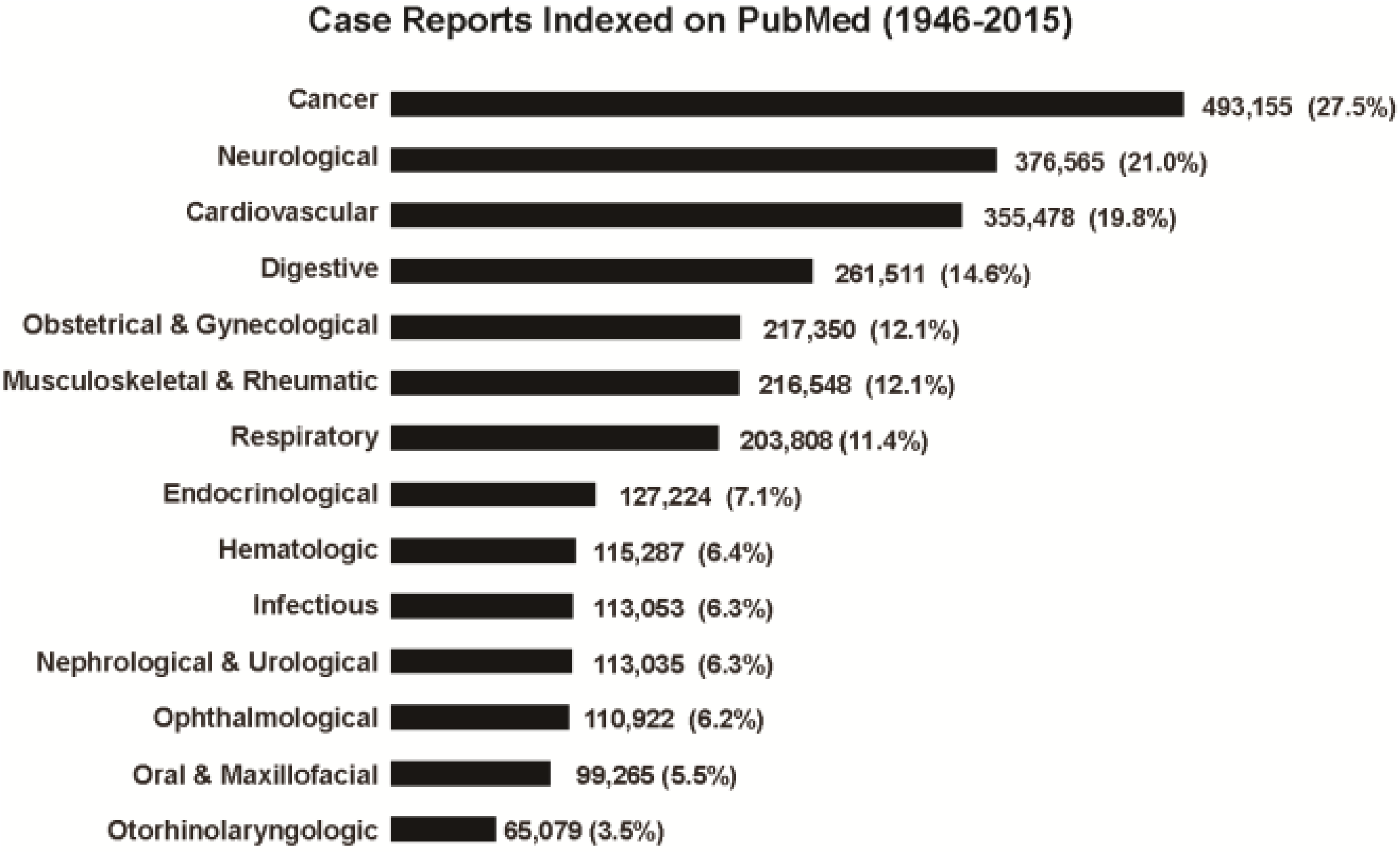
Case Report Publications Per Disease Phenotype. The total number and percentage (in parentheses) of all case reports indexed on PubMed (1946-2015) are displayed for each of fourteen disease phenotypes (by MeSH terms), including cancer, neurological, cardiovascular, digestive, obstetrical & gynecological, respiratory, musculoskeletal & rheumatic, endocrinological, nephrological & urological, hematologic, infectious, ophthalmological, oral & maxillofacial, and otorhinolaryngologic diseases, are shown. Note that the percentages shown do not total to 100% since a case report may belong to more than one disease categories. Cancer, neurological disease, and cardiovascular disease are the three most studied and published areas, accounting for 27.5%, 21.0%, and 19.8% of the total case reports published, respectively.

**Figure 2A.**
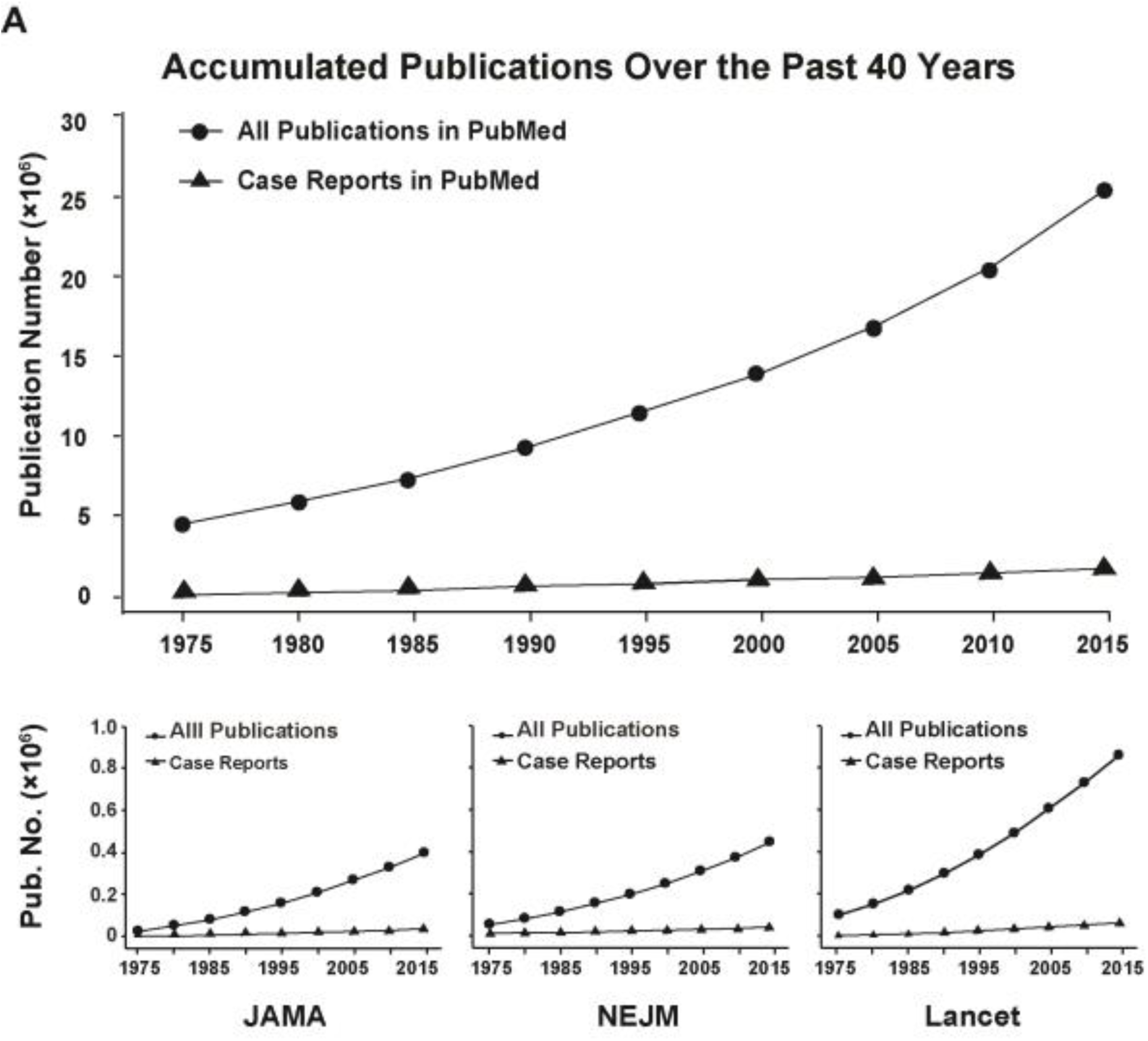
Accumulated Case Report Publications Over Past 40 Years. Top: The accumulated number of publications (X-axis) has been gradually increasing over the last 40 years (1975-2015, Y-axis), though the growth rate of case reports (triangle) has been slower compared to that of all accumulated publications (circle). This may be partially due to the scarcity of rare disease cases and limited citation frequency of case reports, making this form of publication less popular for the journal. Bottom: Examples are shown for case report publications by three prominent journals: NEJM, Lancet, and JAMA. The accumulated number of case report publications (circle) has been likewise increasing at a slower rate compared to all publications (triangle) by these three journals respectively (1975-2015, Y-axis). Note: As PubMed did not annotate many case reports with the article type “Case Reports” until 1975, the displayed number of case reports published before 1975 (∼107,000) may be underestimated and are therefore not shown. Abbreviations: NEJM, The New England Journal of Medicine; JAMA, Journal of American Medical Association.

Recently, the scientific significance of case reports has become a matter of debate. On one side, experts assert that this publication type is undervalued, as the novel findings contained in a case report or case series can catalyze new research such as clinical trials.[10] More importantly, case reports are broadly recognized for serving as a teaching tool by presenting a coherent medical narrative that evolves over time, allowing for reflection on the disease course as well as the reasoning employed to reach a diagnosis and management decision.

Nonetheless, there are also clear limitations to case reports, especially toward serving clinical or scientific research objectives by the biomedical community.[11] First, the enormous amount of case report publications have made it both labor-intensive and time-demanding to manually search and/or process them to identify relevant data.[12] More problematically, the majority of case reports are based on a singular observation and therefore lack statistical evidence and quantitative data. In contrast to conventional clinical studies, clinical cases are not selected from representative patient samples and are insufficient for providing information on disease rates, ratios, incidence, or prevalence.[12] Causal relationships cannot be generated from the observations of an individual patient as the events in one case report are often descriptive, and their observations could be merely coincidental.[13] While individual case reports may contain limited value, identifying their intrinsic connections with other case reports and/or with conventional clinical trials could reveal hidden knowledge regarding the disease and relevant treatments. The development of computational tools to better index and curate case reports through their metadata would enable users to uncover mechanistic insights and improve clinical applications.

However, the creation of tools to extract clinical metadata is obstructed by the very nature of the case reports’ contents, which remain very heterogeneous, unstructured, and sometimes fragmented. Not only does each journal have a different emphasis and structure for the clinical data, but each case report within a journal also has a different style of presentation depending on the patient symptoms and disease progression. For instance, some journals encourage detailed clinical presentations while others prefer comprehensive discussions on differential diagnoses or management decisions. Additionally, each journal may emphasize different types of data (e.g., lab values, images, charts, and links to digital media). Such heterogeneity may pose challenges to efficient extraction of useful information. As a result, little has been accomplished to make this rich source of clinical information discoverable or otherwise useful. Though the National Library of Medicine’s Indexing Initiative and the Semantic Knowledge Representation Project have produced curation and extraction tools like Medical Text Indexer (MTI),[14] SemRep,[15] and MeSHLabeler,[16] these tools primarily function to retrieve keywords, author names, or publication dates.[17] Consequently, development of more efficient curation methods for deeper inquiries will pave the way to the exploitation of accumulated case reports, which currently remain a hidden treasure.

### Standardization of a Metadata Template to Curate Medical Information from Clinical Case Reports

To optimize the utility of clinical case reports in this era of big data and precision medicine, novel algorithms and infrastructure for extracting, indexing, and querying the contents of clinical case reports are urgently needed. Such methods are firmly predicated upon the existence of detailed metadata to accurately describe the contents of case reports. However, metadata in the clinical arena, particularly with respect to case reports, remain poorly defined. Hence, we have carefully analyzed 1000 case reports in PubMed as a test dataset to guide the formulation of a standardized metadata template that reflects the value of case report data and renders case report data more findable, accessible, interoperable, and reusable (FAIR) (Table 1). The template defines 35 key metadata fields; many of them are relevant to the proposed journals’ guidelines for publication of clinical case reports.[18, 19] We have classified the 35 items into three categories. The first category, “Identification”, includes 10 items that serve as identifiers, including title, author, and PMID/DOI numbers. The second category, “Medical Content”, encapsulates 21 items such as disease diagnosis, signs and symptoms, diagnostic procedures, and therapies. Finally, the third category, “Acknowledgements”, contains 4 items regarding external sources of information such as references, funding source, award numbers, and disclosures.

**Table 1.** Standardized Metadata Template for Clinical Case Reports.

| Field | Data Type | Examples (from multiple clinical case reports) |
| --- | --- | --- |
| **Identification (Supports Findability): 10 items** |  |  |
| Title | Textual | Case 3-2017. A 62-Year-Old Man with Cardiac Sarcoidosis and New Diplopia and Weakness. |
| Author | Textual | Samuels MA, R. Gilberto Gonzalez |
| Year | Numerical | 2017 |
| Journal | Textual/Numerical | New England Journal of Medicine |
| Institution | Textual | From the Department of Neurology (M.A.S.), Brigham and Women's Hospital, the Departments of Radiology (R.G.G.), Medicine (A.T.M.), and Pathology (E.T.H.-W.), Massachusetts General Hospital, and the Departments of Neurology (M.A.S.), Radiology (R.G.G.), Medicine (A.T.M.), and Pathology (E.T.H.-W.), Harvard Medical School - all in Boston. |
| Senior Author/PI | Textual | E. Tessa Hedley-Whyte |
| PMID | Numerical | 28121502 |
| DOI | Numerical | 10.1056/NEJMcpc1610713 |
| Link | URL | http://www.nejm.org/doi/full/10.1056/NEJMcpc1610713 |
| Language(s) | Textual | English |
| **Medical content (Supports Findability, Accessibility, Interoperability, Reusability): 21 items** |  |  |
| Keywords | Textual | Sarcoidosis; heart failure; dyspnea |
| Demography | Textual/Numerical | Male/female; age |
| Geographic Locations | Textual | Los Angeles, USA |
| Lifestyle | Textual | Smoker; alcoholic; sedentary; employed |
| Medical History Taking - Family History | Textual | Coronary heart disease; stroke |
| Social Work | Textual | Driver; high school diploma |
| Medical History Taking - Medical/Surgical History | Textual | Graves's disease; Coronary artery bypass surgery |
| Disease System | Textual | Cardiovascular diseases; |
| Signs and Symptoms | Textual/Numerical | Dyspnea; chest pain; rales; edema |
| Comorbidity | Textual | Hypertension; diabetes mellitus |
| Diagnostic Techniques and Procedures | Textual | Echocardiography; electrocardiography |
| Diagnosis | Textual | Acute myocardial infarction; Takotsubo Cardiomyopathy |
| Laboratory Values | Textual/Numerical pairs | Troponin (1.2 ng/mL); hemoglobin (13.1 g/dL) |
| Pathology | Textual | Histology: tumor epithelial cells |
| Drug Therapy | Textual | Metoprolol, aspirin |
| Therapeutics | Textual | Percutaneous coronary intervention; transcatheter aortic |
|  |  | valve replacement |
| Patient Outcome Assessment | Textual | Diseased |
| Diagnostic Imaging/Videotape Recording | Image/Video | https://www.ncbi.nlm.nih.gov/pmc/articles/PMC5223779/figure/f1-amjcaserep-18-7/ |
| Relationship to Other Case Reports | Textual/Numerical | PMID: 24755359. Extra-gastrointestinal stromal tumor of the pancreas: case report and review of the literature. World J Surg Oncol. 2014;12:105. |
| Relationship to Published Clinical Trials | Textual/Numerical | NCT01492725, ACTRN12611000969965 |
| Crosslink with Database | URL | OMIM (Online Mendelian Inheritance in Man) https://www.omim.org/entry/192600 |
| **Acknowledgements: 4 Items** |  |  |
| Funding Source | Textual | NIH |
| Award Number | Textual/Numerical | R01HL123060 |
| Disclosures/Conflict of Interest | Textual | Medtronic, Abiomed, St. Jude; none |
| References | Numerical | 18 (e.g., Sood N, Taub C. Unicuspid aortic valve: An interesting presentation. Eur Heart J. 2008;29:1295.) |

Note: A standardized metadata template outlining 35 essential items that adequately reflect the necessary information. These 35 items are categorized as case report “Identification”, “Medical Content”, and “Acknowledgements”. The expected data types and examples for each item are provided.

To assess the quality and value of the case report data, we devised and applied metrics based on the metadata template to the 1000 case reports. In our scoring system, we assigned one point to each item, for a total possible score of 35. We defined this optimal score of 35 as 100%; likewise, 10 is the highest score for the “Identification” category, 21 for the “Medical Content”, and 4 for the “Acknowledgements” (Figure 3). To determine whether the PubMed metadata sufficiently portrays the contents of the case reports, we scored the metadata manually extracted from case report contents and compared that with the metadata currently available on PubMed for each case report. Our analysis revealed that, overall, case report contents contained an average of 68.4±0.3% of the 35 items, which is significantly higher compared to the existing metadata on published case reports, which contained 47.1±0.2% of the 35 items. Our findings inform that the existing metadata of case reports is often inadequate, and more inclusive metadata are required to better represent key clinical information in case reports, thereby enabling more accurate search results. Future investigations applying autonomous curation of key information via machine learning approaches, such as natural language processing and information retrieval, will be key to enriching metadata for the massive body of case reports. Moreover, search algorithms to identify similar patterns of clinical demographics, pathogenesis, and therapeutic approaches could elevate the interoperability and reusability of case report data.

**Figure 3:**
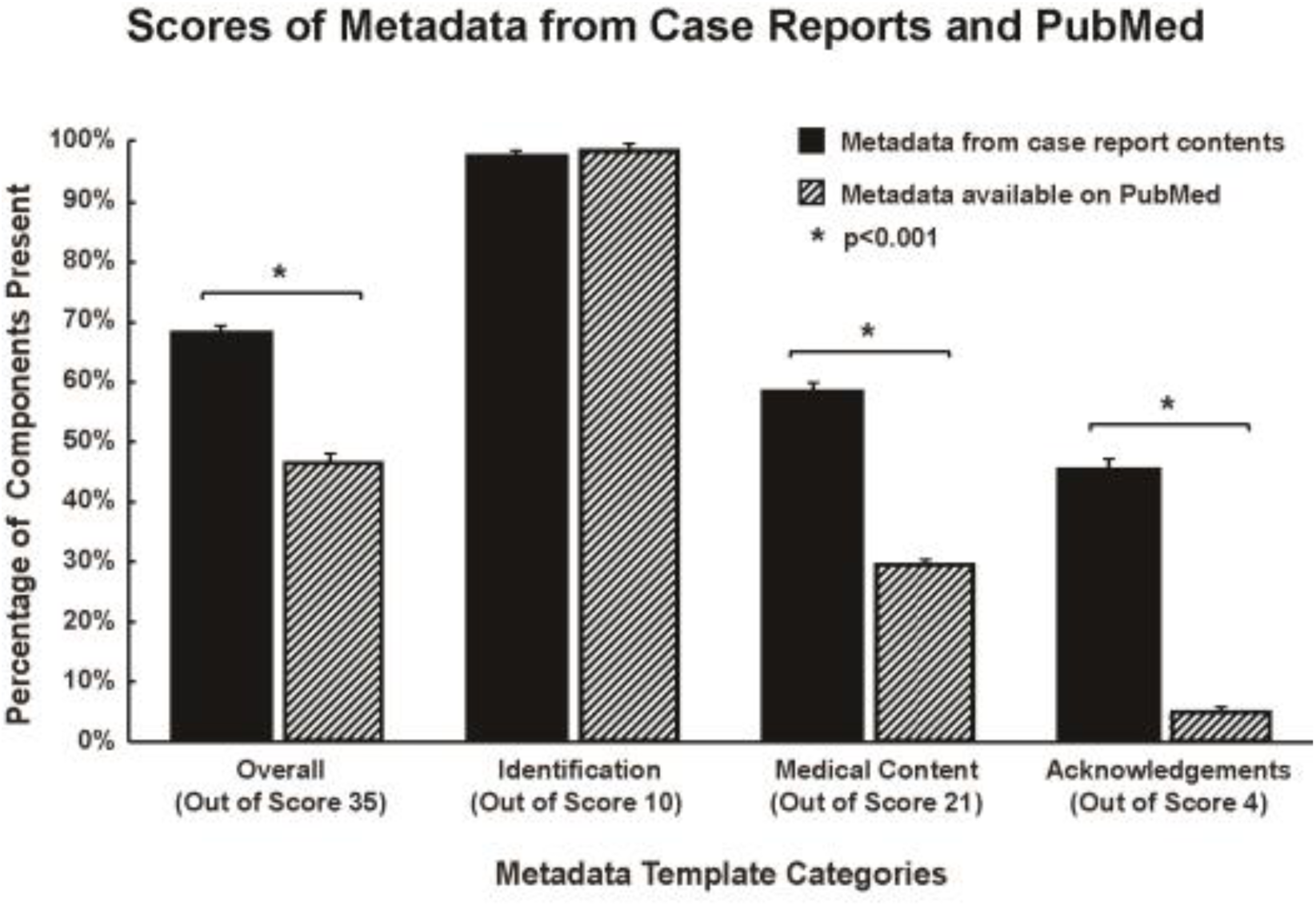
Assessment of Quality of Case Report Metadata. We designed metrics based on the metadata template (Table 1) to assess the quality of the existing metadata for 1000 case reports in PubMed, as well as the value of additional metadata that could be retrieved from the published reports. We defined the optimal score of 35 as 100%, among which 10 is the highest score for the “Identification” category, 21 for the “Medical Content”, and 4 for the “Acknowledgements”. The X-axis displays the metadata template categories and their respective optimal scores. The Y-axis displays the actual percentage of metadata items present in the case reports. We scored the metadata from case report contents (black) and the metadata available on PubMed (striated) for each of the 100 case reports and compared the average scores (as percentages) for each metadata template category. Overall, metadata from case report contents contained an average of 68.4±0.3% of the 35 items, which significantly differs from the metadata available on PubMed, which contained 47.1±0.2% of the 35 items. Our findings inform that the existing metadata of case reports is often inadequate, and more inclusive metadata are required to better represent key clinical information in case reports. (*p<0.001 for Wilcoxon signed-rank test)

### Computational tools to retrieve and analyze clinical case reports

To address how frequently case reports are cited, we designed, created, and implemented a Python-supported algorithm (available on https://github.com/bleakley/clinical-case-reports). This program enabled us to analyze 1,374,513 English-language case reports indexed in PubMed in the past 70 years (1945-2015) with regard to their citation count by PubMed Central articles. Nearly 95% of the case reports are rarely cited over 70 years. Indeed, 54.0% have never been cited and 41.7% have been cited one to five times (Figure 2B). Nonetheless, as many as 23,216 case reports (1.7%) received more than 10 citations, demonstrating the role of case reports as sources of biomedical knowledge. A previous study reported <1% of case reports were cited more than 10 times; however, as this study investigated publications and citations over two sets of 2-year periods (1991-1993 and 2001-2003), their results may not be directly comparable to our findings.[20]

**Figure 2B.**
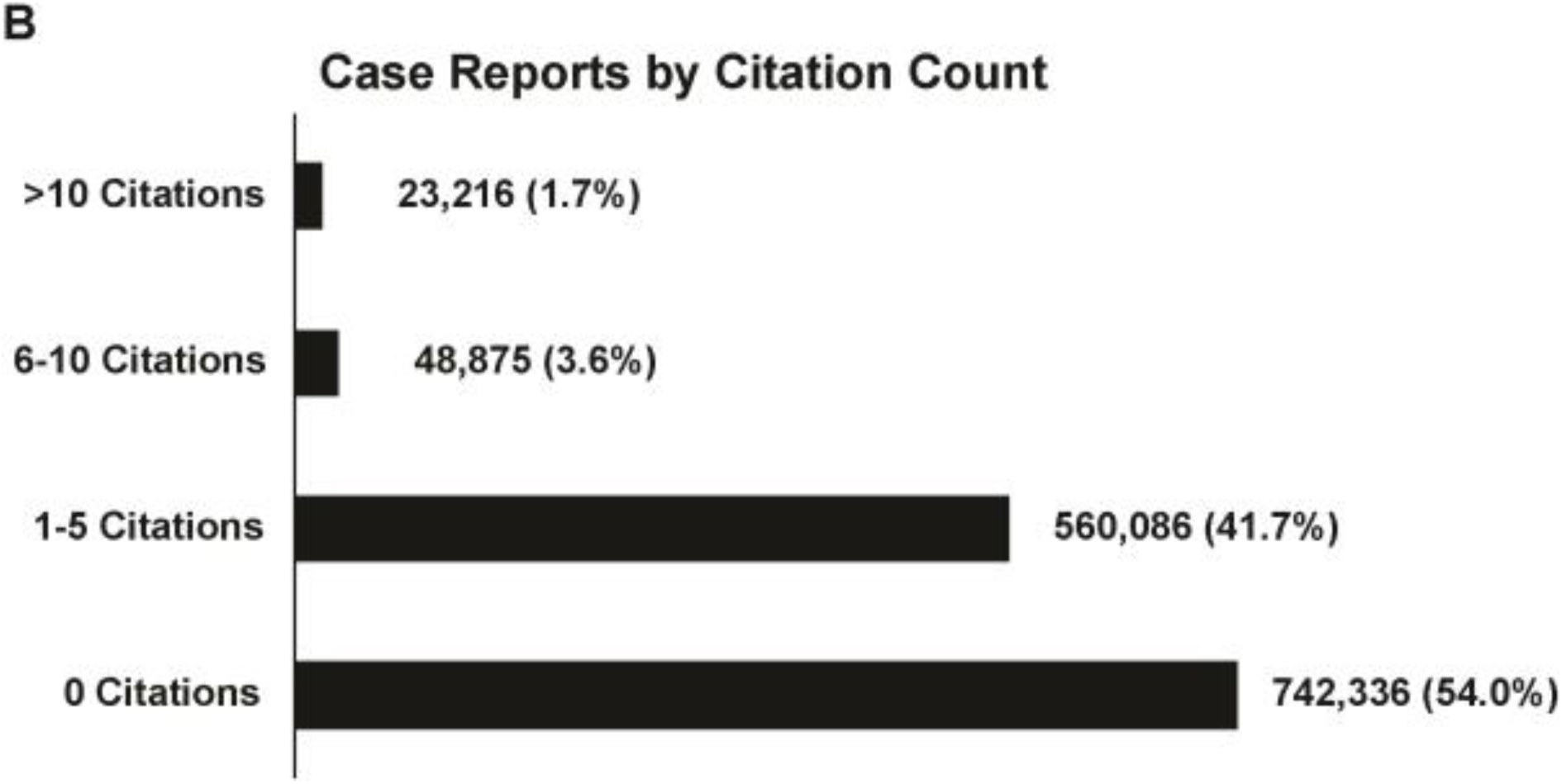
Citations of Case Reports by PubMed Central Articles. The distribution of case reports according to citation frequency by PubMed Central articles is illustrated as number and percentage of all case reports. Citation frequency was classified into categories of 0 citations, 1-5 citations, 6-10 citations, and >10 citations. The majority of case reports indexed in PubMed Central have been poorly cited, with 54.0% that have never been cited and 41.7% that have been cited less than 5 times. Nonetheless, as many as 23,216 case reports (1.7%) received more than 10 citations, demonstrating the role of case reports as sources of biomedical knowledge.

### Broadening clinical case report readership and impact

While, at present, the majority of case reports are read by academic physicians for educational purposes, implementation of the standardized metadata template to make metadata more FAIR can expand the audience and application of case reports. For example, case report user groups may include medical students, interns and fellows, epidemiologists, and statisticians, who may derive valuable information from improved indexing and categorization of case report metadata to better understand clinical phenotypes or to draw relationships of an individual case to a larger representative patient population. As another example, healthcare organizations and policymakers (e.g., FDA) can retrieve case report (meta)data as an additional source for tracking unusual disease occurrences, epidemiological trends, and post-marketing drug surveillance. Moreover, pharmaceutical industries can design a survey on case reports of drugs with unexpected indications or unnoticed side-effects to assist in modifying usage instructions and directing future development.

To address the key metadata items commonly missing in case reports, we envision a solution that integrates what PubMed has already accomplished with MeSH terms with additional classification systems such as the International Classification of Diseases (ICD) and International Classification of Health Interventions (ICHI) to compensate for the missing items when implementing MeSH terms alone. This integration would produce curated, indexed, and structured (meta)data from case reports that can ultimately interface with preclinical omics research, clinical cohort studies, and clinical trials to advance understanding of disease progression, management, and clinical outcome. To surmount the ever-growing amount of free text information with limited annotation and accessibility, computational platforms based on machine learning approaches and in-depth search algorithms will drive better understanding of case reports, clinical trials, and other relevant databases to enable text data FAIR, advance medical science, and improve patient care.

## Conclusion

As a time-honored tradition in medical publication and a treasured source of clinical data, clinical case reports augment our understanding of disease etiology, pathogenesis, miscellaneous diagnosis, and therapeutic effectiveness. The growing volume of case reports published each year stands testament to their popularity and usefulness to their targeted clinical readership, but this size, coupled with the isolated, unstructured, and heterogeneous nature of case reports’ contents, also presents a challenge to index, annotate, and query case report data. In this report, we created a resource platform that enable us to automatically retrieve relevant medical information and to analyze all published PubMed clinical case report. Accordingly, we constructed a standardized metadata template and metrics, as well as a test dataset consisting of 1000 CCRs spanning 14 disease phenotypes, to evaluate the caliber of the existing metadata employed for case reports in PubMed and confirmed a discrepancy between the medical content and the metadata meant to describe it. Our CCR test dataset may be a valuable resource for biomedical researchers who are developing machine learning approaches to advance medical science and improve patient care.

## Contributors

All authors contributed to the writing and final review of the manuscript.

## Acknowledgement

None.

